# Gut microbial metabolite imidazole propionate impairs endothelial cell function and promotes the development of atherosclerosis

**DOI:** 10.1101/2024.06.05.597678

**Authors:** Vanasa Nageswaran, Alba Carreras, Leander Reinshagen, Katharina R. Beck, Jakob Steinfeldt, Marcus Ståhlman, Pegah Ramezani Rad, Joseph Lim, Elisabeth T. Strässler, Barbara Verhaar, Yvonne Döring, Christian Weber, Maximilian König, Elisabeth Steinhagen-Thiessen, Ilja Demuth, Nicolle Kränkel, David M. Leistner, Max Nieuwdorp, Petra Knaus, Marc Ferrell, Michael Potente, Stanley L. Hazen, Ulf Landmesser, Fredrik Bäckhed, Arash Haghikia

## Abstract

**Background:** The microbially generated amino acid-derived metabolite imidazole propionate (ImP) contributes to the pathogenesis of type 2 diabetes. However, the effect of ImP on endothelial cell physiology and its role in atherosclerotic coronary artery disease (CAD) is unknown. Using both human and animal model studies, we investigated the potential contributory role of ImP in the development of atherosclerosis.

**Methods:** Plasma levels of ImP were measured in patients undergoing elective cardiac angiography (n = 831) by means of ultra high-performance liquid chromatography coupled to tandem mass spectrometry. Odds ratios (ORs) and corresponding 95% confidence intervals for CAD were calculated based on the ImP quartiles using both univariable and multivariable logistic regression models. Atheroprone apolipoprotein E^-/-^ (*Apoe^-/-^*) mice fed a high-fat diet were additionally treated with ImP (800 µg) or vehicle and aortic atherosclerotic lesion area was evaluated after 12 weeks. In a mouse model of carotid artery injury, the effect of ImP on vascular regeneration was examined. Using human aortic endothelial cells (HAECs) the effect of ImP on functional properties of endothelial cells were assessed. Next-generation sequencing, western blot analysis, siRNA-based gene knockdown and tamoxifen-inducible Cre-loxP experiments were performed to investigate ImP-mediated molecular mechanisms.

**Results:** Plasma ImP levels in subjects undergoing cardiac evaluation were associated with increased risk for prevalent CAD. In atheroprone *Apoe^-/-^* mice ImP increased atherosclerotic lesion size. We found that ImP dose-dependently impaired migratory and angiogenic properties of human endothelial cells, and promoted an increased inflammatory response. Long-term exposure to ImP impaired the repair potential of the endothelium after an arterial insult. Mechanistically, ImP attenuated insulin receptor signaling by suppressing PI3K/AKT pathway leading to the sustained activation of the forkhead box protein O1 (FOXO1) transcription factor. Genetic inactivation of endothelial FOXO1 signaling in ImP-treated mice enhanced the angiogenic activity and preserved the vascular repair capacity of endothelial cells after carotid injury.

**Conclusions:** Our findings reveal a hitherto unknown role of the microbially produced histidine-derived metabolite ImP in endothelial dysfunction and atherosclerosis, suggesting that ImP metabolism is a potential therapeutic target in atherosclerotic cardiovascular disease.

## INTRODUCTION

Endothelial cell (EC) dysfunction in the arterial vascular tree is the earliest detectable change in the development of atherosclerosis.^1^ Impaired EC function compromises not only the ability to recover arterial integrity after injury but also promotes the adoption of a pro-inflammatory phenotype that attracts monocytes.^2^ In patients with diabetes, EC dysfunction substantially increases the risk of atherosclerosis development. Targeting endothelial dysfunction is thus an important strategy for atherosclerotic cardiovascular disease (ACVD) prevention.^3^

In recent years, functional alterations of the gut microbiome and its metabolic activity have been increasingly acknowledged as causal factors of insulin resistance^4^ and ACVD.^5^ One such example of causal contribution for metaorganismal pathways in ACVD is the metabolite trimethylamine N-oxide (TMAO), which is derived from microbial metabolism of trimethylamine-containing nutrients such as choline and carnitine.^6^ As shown in several mechanistic animal models, TMAO promotes the development of atherosclerosis and increases vascular thrombogenicity^5–7^ with prognostic and clinical implications for patients with atherosclerotic coronary artery disease (CAD)^8–10^ and ischemic stroke.^10^

Metagenomic approaches have linked specific gut microbiota composition with type 2 diabetes, as characterized by a decrease in the abundance of butyrate-producing bacteria and an increase in distinct opportunistic pathogens.^11–13^ This altered ecosystem is associated with altered microbial metabolism and production of metabolites.^14^ Imidazole propionate (ImP), a histidine-derived metabolite, is increased in the portal and peripheral blood of patients with type 2 diabetes compared to normoglycemic controls.^15,16^ Importantly, when administered to mice, ImP caused glucose control impairment^15^ and attenuated the glucose-lowering effect of metformin^17^, demonstrating a causal role for circulatory ImP in the pathogenesis of type 2 diabetes. Furthermore, ImP independently correlates with reduced ejection fraction and heart failure in clinical cohorts.^18^ However, to date, the effect of ImP on endothelial cell function and its role in ACVD is unknown.

In this study, we sought to determine the relation between circulatory ImP levels and CAD risk in humans. We tested the effect of increased blood levels of ImP on the development of atherosclerotic lesions in an apolipoprotein E^-/-^ (*Apoe*^-/-^) mouse model. Furthermore, we examined the effect of ImP on endothelial cell function and how elevated ImP levels impact endothelial repair capacity after vascular injury in mice.

## METHODS

### Materials availability

All data, analytic methods, and study materials are available to other researchers for purposes of reproducing these results or replicating these procedures by reasonable request directed to the corresponding author. Detailed materials have been incorporated in the article and are provided in the Supplementary Tables 1 – 4.

### Human study

Blood samples were obtained from subjects enrolled in the LipidCardio Study (German Clinical Trial Register; drks.de; Identifier: DRKS0002091). The study was approved by the local research ethics committee (approval number: EA1/135/16) and all participants provided written informed consent. Study protocol and design of the LipidCardio study were described previously.^19^

Patients aged 18 years and older undergoing cardiac catheterization at a single large academic center (Deutsches Herzzentrum der Charité, Campus Benjamin Franklin), except those with troponin-positive acute coronary syndromes (ACS), were eligible for inclusion. Cardiac catheterization and coronary angiography were performed according to the standard protocols of the interventional cardiology unit and by discretion of the interventional cardiologist. The interventional cardiologist routinely documented diagnostic findings. Comprehensive angiographic results at the time of enrolment were recorded in the study database.

Blood samples were frozen immediately following collection and between October 2016 and March 2018, a number of 1005 consecutive patients were enrolled of whom 831 blood samples were available for the current study. Co-morbidities and cardiovascular risk factors, including diabetes, arterial hypertension and dyslipidemia were evaluated.

Patient samples and clinical data used for correlating ImP levels with soluble VCAM-1 levels were obtained from the GeneBank at the Cleveland Clinic study: Molecular Determinants of Coronary Artery Disease (ClinicalTrials.gov; Identifier: NCT00590200). GeneBank is a single site sample repository generated from consecutive patients undergoing elective diagnostic coronary angiography or elective cardiac computed tomographic angiography with extensive clinical and laboratory characterization and longitudinal follow-up. Subjects were recruited between 2001 and 2007. Exclusion criteria for GeneBank included patients with a recent myocardial infarction (< 4 weeks) or elevated troponin I (> 0.03 mg/dl) at enrolment. Subjects with atherosclerotic CAD were defined as patients with adjudicated diagnoses of stable or unstable angina, myocardial infarction, history of coronary revascularization or angiographic evidence of ≥ 50% stenosis of at least one major coronary artery at the time of coronary angiography. All clinical study protocols and informed consent for human individuals were approved by the Cleveland Clinic Institutional Review Board. Written informed consent was obtained from all individuals. Soluble VCAM-1 levels were measured by enzyme-linked immunosorbent assay (ELISA) according to the manufacturer’s protocol (Invitrogen, Waltham, MA).

### Blood tests and Imidazole propionate measurement

Laboratory workup was performed for total cholesterol, low-density lipoproteins, triglyceride, creatinine, and full blood count. Blood samples were immediately processed and stored in the Central Biomaterial Bank Charité (ZeBanC) at −80°C for further analyses. Plasma levels of ImP were quantified using ultra high-performance liquid chromatography (UHPLC) coupled to tandem mass spectrometry. Analyses were performed in two different laboratories in Sweden (LipidCardio) and in the USA (GeneBank at the Cleveland Clinic). Both laboratories have used the same internal standard (13C3-ImP) for their sample preparations. The samples from the European cohort were prepared and analyzed based on a previously published method with minor modifications.^15^ Briefly, 25 µL of plasma samples were extracted using 6 volumes of acetonitrile containing 100 nM of internal standard (ImP-13C3, Astra Zeneca, Cambridge, UK) before drying the samples under a flow of nitrogen. Then, the samples were reconstituted with 5% HCl (37%) in 1-butanol, subjected to n-butyl ester derivatization and finally reconstituted in 150 µL water:acetonitrile (9:1). ImP levels were determined using multiple reaction monitoring of the transitions 197/81 and 200/82 for the internal standard.

The samples of the GeneBank study were subjected to the LC-MS/MS analysis on a chromatographic system consisting of two Shimadzu LC-30 AD pumps (Nexera X2), a CTO 20AC oven operating at 30 °C, and a SIL-30 AC-MP autosampler in tandem with 8050 triple quadruple mass spectrometer (Shimadzu Scientific Instruments, Inc., Columbia, MD, USA). The limit of quantification (LOQ) with a signal-to-noise cutoff of 10:1 was 5 nM. Values below the LOQ were reported as half the LOQ value. Three quality control samples were performed with each sample batch and inter-batch variations, expressed as coefficient of variation (CV) were less than 10 %. For data analysis LabSolution software (version 5.89; Shimadzu) was used. Investigators performing analyses were blinded to sample identity other than barcode label.

### Animal models

Male wild type C57BL/6J and apolipoprotein E knockout (*Apoe^-/-^*) mice were purchased from Charles River Laboratories (Germany). Endothelial-specific FOXO1 knockout mice (*Cdh5-CreERT2*(+/–)*/FOXO1^fl/fl^*) on C57BL/6J background and control littermates without Cre recombinase (*Cdh5-CreERT2*(+/+)*/FOXO1^fl/fl^*) were obtained through collaboration.^20^ All mice experiments were performed in our animal facility and were approved by the ethics committee on animal care and use in Berlin, Germany. Mice were maintained under 12 h light cycle and with access to food and water *ad libitum*.

Atherosclerosis was induced in adult (19-20 weeks of age) male *Apoe*^-/-^ mice fed a high fat diet (HFD, crude fat 34%, cholesterol 290 mg/kg; Ssniff, Germany, E15741) for 12 weeks. Simultaneously, ImP (800 µg/day) (Santa Cruz) or vehicle in drinking water was administrated to *Apoe*^-/-^ mice. All mice were sacrificed after 12 weeks of treatment (Fig. 2A). Blood was collected by cardiac puncture and organs (heart and aorta) were isolated and snap-frozen in liquid nitrogen or dry ice and stored at -80°C for further analysis.

To investigate the effect of ImP on arterial injury, adult (10 weeks of age) male C57BL/6J mice were randomly assigned to either control group (n = 6) or ImP group (n = 6) for a total of 24 days (Fig. 4A). Carotid artery injury in mice was performed as described.^21^ Briefly, mice were first anesthetized with 3% of isoflurane and then maintained with 1.5% isoflurane in oxygen (1l/min) via inhalation and body temperature was maintained using a heating pad kept at 37°C preventing hypothermia. After shaving the left side of the neck, a small incision was made to expose the left carotid artery (LCA). Carotid injury was induced to LCA by an electric impulse of 2 W for 2 seconds over a length of 4 mm using bipolar microforceps (VIO 50 C, Erbe Elektromedizin GmbH). Three days after carotid injury, re-endothelialization was assessed by injecting 50 µl of 5% Evans blue (Sigma) into the left heart ventricle two minutes before sacrifice. LCA was isolated, rinsed in PBS and fixed on microscope slides. The Evans blue-stained denuded area was captured *en face* by a brightfield microscope (Axioskop 40, Zeiss), and re-endothelialization was calculated longitudinally as the ratio of blue-stained area to injured area and subtracted from 100% using imaging software (ImageJ, NIH).

To assess the regulatory role of FOXO1 on vascular regeneration *in vivo*, adult endothelial-specific FOXO1 knockout mice (13 weeks of age) were generated by crossing *loxP*-flanked FOXO1 mice (*FOXO1^fl/fl^*) with transgenic mice expressing the tamoxifen-inducible vascular endothelial-specific cadherin (*Cdh5*) promoter-driven CreERT2 recombinase.^20,22^ Specifically, FOXO1 deletion (*Cdh5-CreERT2*(+/–)*/FOXO1^fl/fl^*) was induced by intraperitoneal injections of tamoxifen (100 mg per kg body weight; Sigma) dissolved in peanut oil (Sigma) and administered once daily for a total of 5 consecutive days. Littermate animals homozygous for the floxed *FOXO1^fl/fl^*but lacking the expression of Cre recombinase were used as control animals (*Cdh5-CreERT2*(+/+)*/FOXO1^fl/fl^*) (Fig. 6C). After tamoxifen induction animals were treated with ImP via drinking water for a total of 3 weeks prior carotid artery injury.

### Lipid analysis

Mouse plasma samples were subjected to fast-performance liquid chromatography (gel filtration on Superose 6 column (GE Healthcare). Different lipoprotein fractions were separated and evaluated based on flow-through time. Cholesterol levels were quantified using an enzymatic assay (Cobas, Roche) according to the manufacturer’s protocol.

### *En face* determination of atherosclerosis

Atherosclerotic plaques were visualized using an *en face* method. Aortic arch and distributing branches were dissected from the mouse and opened longitudinally using a stereomicroscope (SMZ745T, Nikon). Following overnight fixation with 4% paraformaldehyde (PFA), aortas were stained with Oil Red O (ORO) and aortic arch plaque area was calculated as percentage of ORO-stained area to total aortic arch area.

### Histology

For histological analysis of the aortic roots at the base of the heart (basis cordis), tissues were embedded in Tissue-Tek OCT (Sakura) and frozen at −80°C until further processing. 6 µm cryo-sections of the aortic root were prepared and stained for ORO to evaluate atherosclerotic plaque formation. Hematoxylin/eosin (HE) staining was used to assess aortic root morphology. The percentage plaque area to total aortic root area was calculated using ImageJ software.

### Immunohistochemistry

Immunohistochemistry of frozen sections of the aortic roots was performed using the Avidin-Biotin Complex (ABC) kits (Vector Laboratories). Briefly, acetone-fixed sections were washed in 1x PBS (without Ca^2+^ and Mg^2+^) before treatment with 0.075% H_2_O_2_ for 10 min to block endogenous peroxidase activity. The slides were then blocked with 10% rabbit serum (Dako) in Avidin-Biotin blocking solution (Vector Laboratories) for 30 min each. Sections were incubated with primary antibody against CD68 (Abcam, 1:100) for 1 h and washed in PBS before being incubated with anti-rat biotinylated secondary antibody (Dako, 1:100) for 1 h. The slides were then treated with an ABC-HRP Kit for Peroxidase (Vector Laboratories) for 30 min followed by AEC (3-amino-9-ethyl-carbazole, Sigma) staining for 12 min. Aortic roots were counter-stained in hematoxylin solution (Carl Roth) for 15 s and washed in deionized water before mounting with gelatin medium. Macrophage accumulation was determined in plaque area by calculating the percentage of CD68-positive staining to total aortic root area.

### Cell culture

Primary human aortic endothelial cells (HAECs) (Cell Applications; Table S1) were used between passages 7 and 9 for experiments. HAECs were cultured in Endothelial Growth Medium-2 (PromoCell GmbH) supplemented with 10% fetal bovine serum (FBS), 100 units/ml penicillin and 100 µg/ml streptomycin. Human THP-1 monocytes were cultured in RPMI 1640 medium (Gibco) supplemented with 10% FBS, 2 mM L-glutamine, 100 U/mL penicillin and 100 μg/mL streptomycin. Cells were grown to confluence at 37°C and 5% CO_2_ prior to experiments.

### Wound healing assay

Endothelial wound healing potential was assessed by cell scratch assay *in vitro*. HAECs (6 x 10^4^ /well) were seeded on fibronectin-coated 24-well culture plate overnight, and then serum-starved (0.5% FBS) for 5 h at 37°C in cell culture incubator prior to stimulation. The cells were scratched using a sterile yellow pipette tip and treated with 10 nM ImP, 100 nM ImP, 10 ng/ml recombinant TNF-α or control for 16 h. Pictures of the wound area were taken at times 0 h and 16 h post migration using a phase-contrast microscope (EVOS XL Core; Thermo Fisher Scientific). The width of the scratch was measured and quantified by NIH ImageJ software.

### Tube formation assay

To evaluate the effect of ImP on tube formation, 1.5 x 10^4^ HAECs/well were cultured on Matrigel using growth factor-reduced basement membrane extract (BME; Gibco) in 96-well flat bottom plate and stimulated in conditioned medium with control, 50 ng/ml IGF-1, 100 nM ImP or 10 ng/ml recombinant TNF-α for 16 h. After tubular network was formed by HAECs, cells were stained with 6 µM Calcein AM solution (R&D Systems) for 15 min at 37°C. Representative images were taken using an inverted fluorescent microscope (BZ-X; Keyence Corporation), and tube formation was evaluated with „Angiogenesis Analyzer” tool in ImageJ software by quantifying the total number of segments.

### Flow cytometry of HAECs

Expression of cellular adhesion molecules of ICAM-1, VCAM-1 and E-selectin were assessed by fluorescence-activated flow cytometry (FACS). HAECs (6 x 10^4^ /well) were seeded on 24-well plates and stimulated with control medium, 10 nM ImP or 100 nM ImP for 24 h before being washed and collected into FACS tubes (BD Biosciences). Cells were incubated with following antibodies: CD54 (AF700, HA58, 1:100), CD106 (APC, STA, 1:100) and CD62E (PE/Cy7, HAE-1f, 1:100) (Table S2) for 15 min at room temperature. All antibodies were purchased from BioLegend. Fluorescence-minus controls (FMOs) and unstained HAECs were included in the measurements. Samples were acquired using Attune Nxt Flow Cytometer (Thermo Fisher Scientific) and cells were analyzed with Kaluza software (Beckman & Coulter).

### Flow adhesion assay

Monocyte-endothelial cell interaction was investigated by a flow-based adhesion assay. HAECs were seeded in ibidi y-shaped chamber slides (ibidi GmbH) for 5 h and then incubated overnight under sterile flow conditions using a shear stress of 20 dyn/cm^2^. After acclimatization, cells were treated with control, 10 nM ImP or 100 nM ImP for 24 h, respectively. Finally, THP-1 monocytes (1 × 10^6^ cells/ml) were labeled with DiI cell staining solution (Invitrogen) before being perfused through the chambers for 30 min. After incubation, non-adherent monocytes were removed with PBS and co-cultures were fixed with 4% PFA. The number of adherent monocytes to HAECs was quantified from different fields of view using a fluorescence phase-contrast microscope (BZ-X; Keyence Corporation).

### Protein extraction and Western blotting

Whole-cell lysates from washed endothelial cells were obtained using RIPA buffer (50 mM Tris, 150 mM NaCl, 1 mM EDTA, 1 mM NaF, 1 mM DTT, 10 mg/mL aprotinin, 10 mg/mL leupeptin, 0.1 mM Na_3_VO_4_, 1 mM PMSF, and 0.5% NP-40, pH 7.5) (Table S4). Lysates were centrifuged for 10 min at 14.000 rpm and 4°C followed by protein determination with Pierce^TM^ BCA Protein Assay Kit (Thermo Scientific) according to manufacture description. 20 ug of proteins were separated by 10% SDS-PAGE and transferred onto a PVDF membrane (Merck). Membranes were blocked in 5% milk or bovine serum albumin (BSA) for 1 h at room temperature and incubated with primary antibodies against: PI3 Kinase Class II α (Cell Signaling, 1:2000), Phospho-AKT(Ser473) (Cell Signaling, 1:1000), AKT (Cell Signaling, 1:1000), Phospho-FOXO1(Thr24) (Cell Signaling, 1:2000), FOXO1 (Cell Signaling, 1:2000) and GAPDH (Merck, 1:10–000) (Table S2) overnight at 4°C. The membranes were then incubated with horseradish-peroxidase-conjugated anti-rabbit or anti-mouse secondary antibody (Southern Biotechnology) for 1 hour at room temperature. Proteins were detected with SuperSignal® West Dura Extended Duration Substrate (Thermo Scientific) and the intensity of chemiluminescence was measured using a UVP ChemStudio PLUS imaging system (Analytik Jena).

### Immunocytochemistry

HAECs were grown on fibronectin-coated glass slides and treated with versus without ImP (100 nM) for 24 h. Cells were then washed and fixed with 4% PFA for 15 min at room temperature. Permeabilization was performed in 0.5% Triton X-100 in PBS for 15 min and blocking in 5% horse serum for 1 h. HAECs were stained overnight for anti-FOXO1 (Cell Signaling, 1:100) and DAPI (Roche Diagnostics, 0.1 µg/ml) followed by DyLight 594-conjugated goat anti-rabbit secondary antibody (Invitrogen, 1:100) (Table S2) for 1h. Coverslips were washed and mounted in Kaiser′s glycerol gelatin (Merck). The fluorescence intensity of nuclear FOXO1 staining was analyzed with a Keyence fluorescence microscope.

### siRNA transfection

HAECs were plated at a density of 3 x 10^5^ cells/well in 6-well format before being transfected with FOXO1 small interfering RNA (siRNA) or scramble siRNA (ON-TARGETplus, Dharmacon) (Table S3) for 4 h in Opti-MEM medium (Gibco) using Lipofectamine RNAiMAX transfection kit (Invitrogen) according to the manufacturer’s protocol. FOXO1 gene silencing efficacy was confirmed on protein levels by Western blot analysis.

### RNA sequencing

Total RNA was extracted from endothelial cells after stimulation with ImP (100 nM) or control for 12 h using the RNeasy Mini Kit (Qiagen) according to manufacturer instructions. RNA purity and yield were assessed with NanoVue (GE Healthcare), and ribosomal RNA integrity was confirmed by agarose gel electrophoresis stained with ethidium bromide. Three replicates from three independent experiments were carried out for next generation sequencing (NGS). RNA library preparation, sequencing and alignment of the reads was outsourced to GENEWIZ from Azenta Life Sciences (Leipzig, Germany). Library preparation was performed using the NEBNext Ultra II Directional RNA Library Prep Kit following the manufacturer’s instructions (oligodT enrichment method). The samples were sequenced on an Illumina Novoseq 6000 platform with 150bp paired-end reads. Sequences were trimmed to remove adapter sequences and nucleotides with poor quality using Trimmomatic (v0.36). The trimmed reads were mapped to the human genome reference ENSEMBL version 86 using the STAR aligner (v2.5.2b). Gene counts were calculated with the featureCounts program in the Subread package (v1.5.2), only counting unique gene hits. The mean number of total reads per sample was 29.2±3.5 million, of which 99.6% were mapped, resulting in on average 25±3.8 million counts per sample.

### Statistical analysis

Statistical analyses for the *in vitro* and *in vivo* analyses were performed by GraphPad Prism 9 (GraphPad Software, Inc) and presented as mean ± s.e.m. Normal distribution of data was assessed using the Shapiro-Wilk test and was analyzed by unpaired two-tailed Student’s *t*-test (for two groups) or one-way ANOVA followed by Bonferroni’s post hoc test for multiple comparisons (for ≥ 3 groups). Otherwise, nonparametric statistical analyses were performed using the Mann-Whitney *U*-test for two groups or the Kruskal-Wallis test followed by the Dunn post hoc test for multiple comparisons (for ≥ 3 groups).

Differential gene expression (DGE) was tested using the DESeq2 package (v1.36.0) in RStudio (v2022.07.2+576) with R (v4.2.1) ^23^. For these analyses, the formula Gene ∼ Condition was used, with a Wald test and parametric fit. *P*-values were adjusted using false discovery rate (FDR) multiple testing correction. Genes with an adjusted *P*-value < 0.1 were considered as differentially expressed between control and ImP. Gene counts were normalized using DESeq2’s median of ratios method to account for sequencing depth and RNA composition ^23^. A volcano plot was drawn using ggplot2 (v3.4.0). After zero mean, unit variance scaling of the normalized gene counts, a heatmap with differentially expressed genes was drawn using the ComplexHeatmap package (v2.12.1). The dendrogram in this plot was constructed with Ward’s method. The code for the DGE analyses was shared in a Github repository (https://github.com/barbarahelena/transcriptomics-imp-charite).

In human study odds ratios (ORs) and corresponding 95% confidence intervals (CIs) for CAD were calculated based on the ImP quartiles using both univariable and multivariable logistic regression models. In the multivariable models, adjustments were made for age and sex only, as well as established cardiovascular risk factors including age, sex, smoking status, hypertension, hypercholesterolemia, and diabetes. Significance was assumed at a two-sided *P*-value of *P* ≤ 0.05.

### Data availability

RNA sequencing data were made publicly available in the European Nucleotide Archive (ENA) under accession number PRJEB61803 (https://www.ebi.ac.uk/ena/browser/view/PRJEB61803).

## RESULTS

### Plasma levels of imidazole propionate are associated with CAD

To investigate whether ImP is associated with CAD, we examined ImP plasma levels and CAD risks in a patient cohort (n = 831) undergoing elective cardiac evaluation. Patient demographics, laboratory values, and clinical characteristics of all patients are provided in Table 1. We also compared these parameters between patients with and without CAD. Plasma levels of ImP were increased in patients with significant (≥ 50% stenosis) angiographic evidence of CAD, as revealed by diagnostic cardiac catheterization (Figure 1A) with a dose-dependent relationship between ImP levels and risk of prevalent CAD (Figure 1B). This increased risk was particularly pronounced in patients in the fourth quartile and remained significant after the adjustments for both age and sex (Model 1 OR: 3.80; 95% CI, 2.39 - 6.15), and even after further adjustments for cardiovascular risk factors including hypertension, dyslipidemia and smoking (Model 2 OR: 4.22; 95% CI, 2.60 - 6.97). Notably, we found significantly higher levels of ImP in patients with diabetes than in controls (Figure S1), confirming previous findings demonstrating diabetogenic effects of ImP.^15^

**Table 1:**
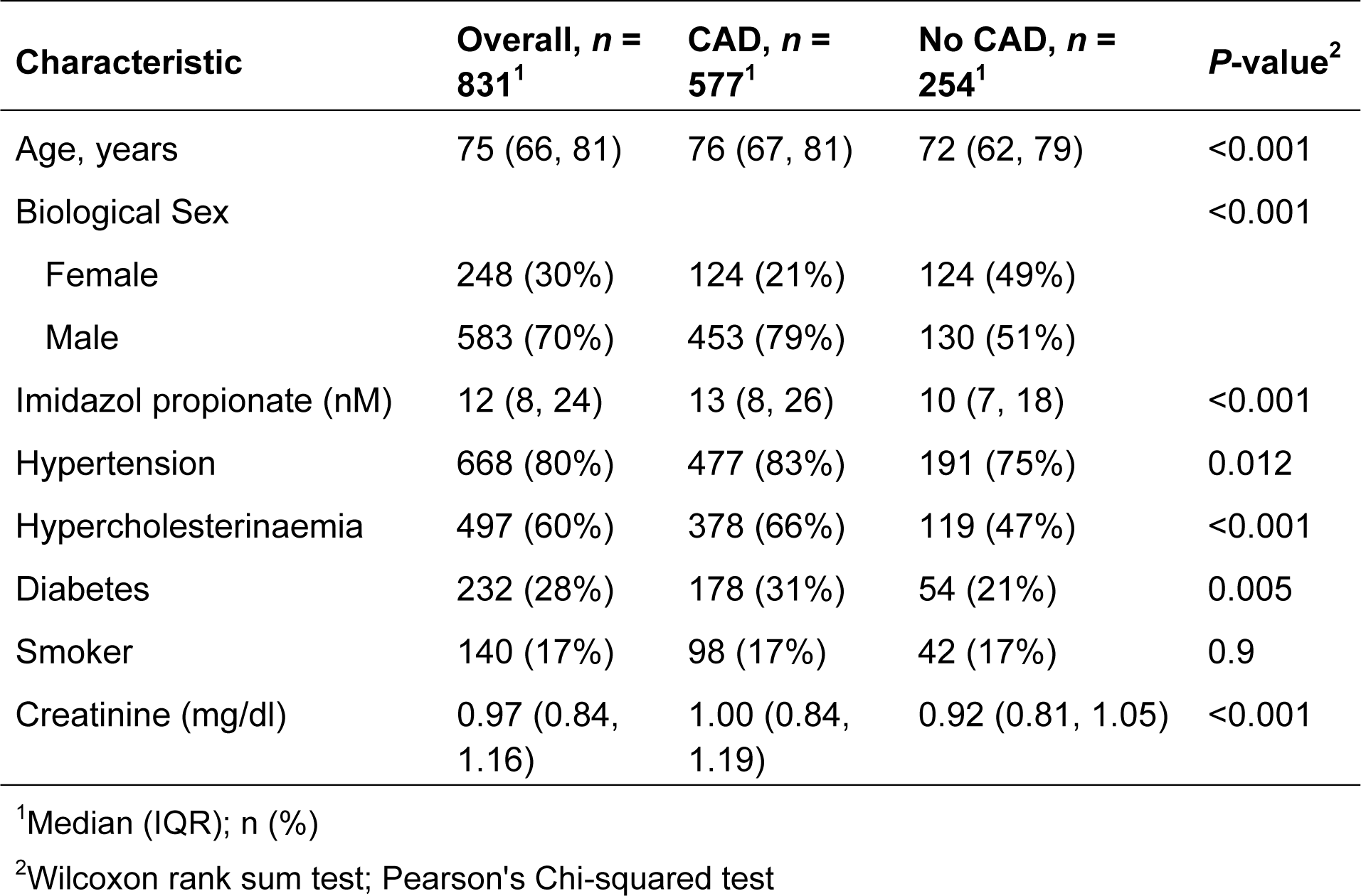
Baseline clinical parameters and laboratory test results.

| Characteristic | Overall, *n* = 831<sup>1</sup> | CAD, *n* = 577<sup>1</sup> | No CAD, *n* = 254<sup>1</sup> | *P*-value<sup>2</sup> |
| --- | --- | --- | --- | --- |
| Age, years | 75 (66, 81) | 76 (67, 81) | 72 (62, 79) | <0.001 |
| Biological Sex |  |  |  | <0.001 |
| Female | 248 (30%) | 124 (21%) | 124 (49%) |  |
| Male | 583 (70%) | 453 (79%) | 130 (51%) |  |
| Imidazol propionate (nM) | 12 (8, 24) | 13 (8, 26) | 10 (7, 18) | <0.001 |
| Hypertension | 668 (80%) | 477 (83%) | 191 (75%) | 0.012 |
| Hypercholesterinaemia | 497 (60%) | 378 (66%) | 119 (47%) | <0.001 |
| Diabetes | 232 (28%) | 178 (31%) | 54 (21%) | 0.005 |
| Smoker | 140 (17%) | 98 (17%) | 42 (17%) | 0.9 |
| Creatinine (mg/dl) | 0.97 (0.84, 1.16) | 1.00 (0.84, 1.19) | 0.92 (0.81, 1.05) | <0.001 |

<sup>1</sup>Median (IQR); *n* (%)
<sup>2</sup>Wilcoxon rank sum test; Pearson's Chi-squared test

**Figure 1:**
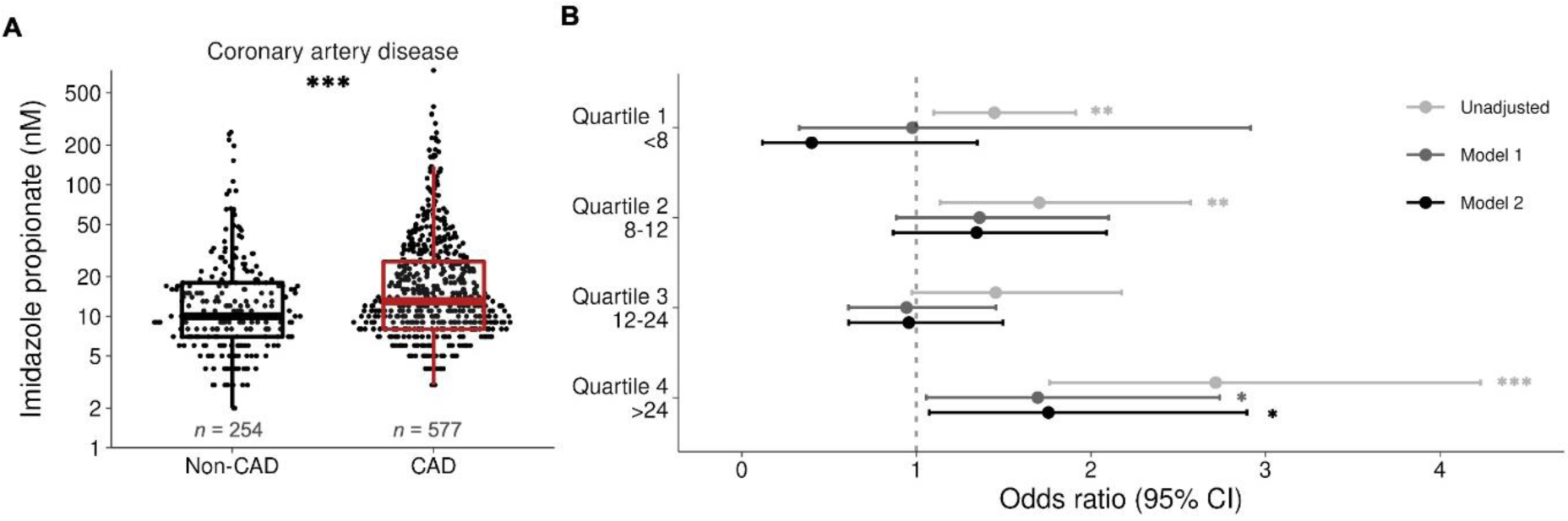
Increased ImP plasma levels are associated with atherosclerotic cardiovascular disease. **A,** Box-whisker plots of circulating ImP levels in patients without coronary artery disease (Non-CAD) as compared to patients with CAD. Data is represented as boxplots: the middle line is the median, the lower and upper hinges are the first and third quartiles, and the whiskers represent 10th and 90th percentiles. **B,** Risk of CAD among all test subjects according to ImP quartile levels using multivariable logistic regression models. Unadjusted odds ratio, adjusted Model 1 (age, sex) and adjusted Model 2 (Model 1 plus incidence of hypertension, dyslipidemia, diabetes and smoking). Symbols represent odds ratio, and the 95% confidence intervals (CI) are indicated by the line length. Data are shown as mean ± SEM and were calculated using Wilcoxon rank sum test (*n* = 831). *\*P* < 0.05, *\*\*P* < 0.01, *\*\*\*P* < 0.001.

### Imidazole propionate aggravates High Fat Diet-induced atherosclerosis in *Apoe*-/-mice

To investigate if ImP could directly promote CAD, we assessed atherosclerosis development in *Apoe*^-/-^ mice receiving ImP via drinking water for 12 weeks (Figure 2A). As expected, this protocol increased plasma ImP levels 12-fold (Figure 1D), which is in range with the patients with highest levels of ImP (Figure 2B). Treatment with ImP did not affect blood levels of total cholesterol (TC), nor of the atheroprone very low-density lipoprotein (VLDL) and low-density lipoprotein (LDL) cholesterol compared to the control group (Figures 2C and 2D). In contrast, chronic ImP administration increased atherosclerotic lesion size after 12 weeks with increased accumulation of CD68^+^ macrophages in the atherosclerotic lesion area (Figures 2E and 2F), highlighting pro-atherogenic and pro-inflammatory effects of ImP, independent of blood lipoprotein levels.

**Figure 2:**
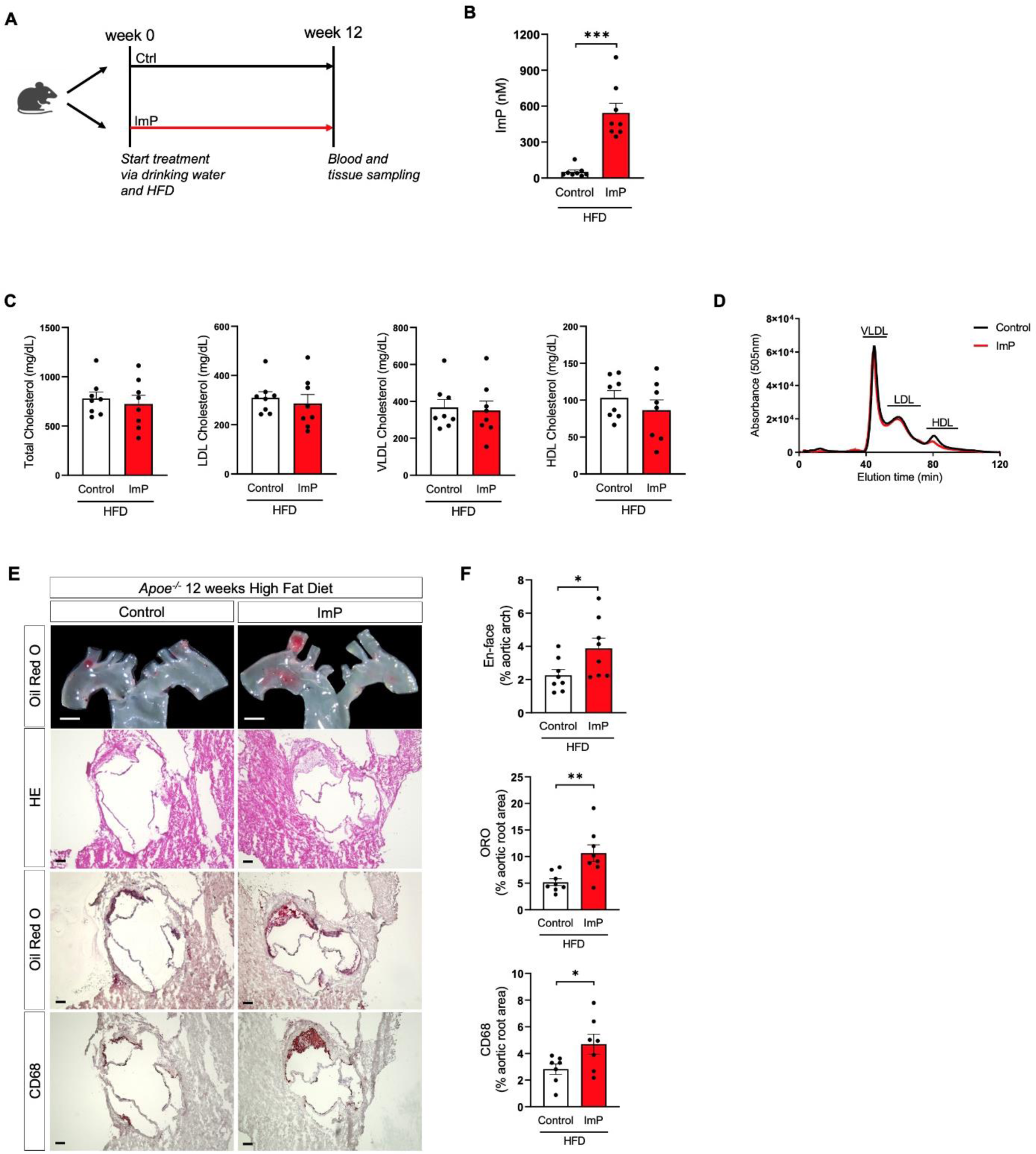
ImP aggravates HFD-induced atherosclerosis in *Apoe-/-* mice. **A,** Schematic illustration of the mouse model of atherosclerosis. *Apoe^-/-^* mice were fed a high-fat diet (HFD) and treated with versus without ImP (800 µg/day) in drinking water for 12 weeks (*n* = 8). **B,** Plasma levels of ImP in *Apoe^-/-^* mice after 12 weeks treatment of HFD and exposure to vehicle or ImP (800 µg/day). **C,** Evaluation of levels of total cholesterol (TC), very low-density lipoprotein (VLDL), LDL cholesterol and HDL cholesterol in control mice versus ImP-treated mice fed a HFD. **D,** HPLC-derived lipoprotein fractions in plasma from control versus ImP mice fed with HFD. Lipoproteins were detected by absorbance at 505 nm and at different elution times. **E,** Representative images of Oil Red O (ORO)-stained *en face* aortic arch and of aortic root sections stained with hematoxylin/eosin (HE), ORO and CD68 (scale bars represent 1mm for *en face* images or 100 µm for aortic root images). **F,** Quantification of atherosclerotic plaque area in *en face* aortic arch and of total aortic root area of control versus ImP-treated mice as percentages. Data are shown as mean ± SEM and were calculated by two-tailed Mann-Whitney *U*-test (B) or unpaired two-tailed Student’s *t*-test (C and F) (*n* = 8 per group). *\*P* < 0.05, *\*\*P* < 0.01, *\*\*\*P* < 0.001.

### Imidazole propionate impairs migratory and angiogenic properties of endothelial cells

Given the central role of endothelial dysfunction in atherosclerosis,^24^ we hypothesized that elevated ImP may contribute to the pathogenesis of ACVD by compromising the properties of ECs. We thus treated human aortic endothelial cells (HAECs) with ImP to assess its impact on the migration and angiogenic properties of ECs. ImP significantly reduced gap closure in an *in vitro* scratch injury assay, suggesting an impaired EC migratory capacity (Figures 3A and 3B). To further address if the reduced migratory capacity resulted in defective angiogenic activity, we next used a Matrigel angiogenesis assay. We observed significantly reduced tube formation in ImP-pretreated ECs compared to control cells, indicating that ImP impairs angiogenic potential in ECs (Figures 3C and 3D). Together, these results suggest that ImP may harm EC functions, which are important for wound healing in the vessel wall.

**Figure 3:**
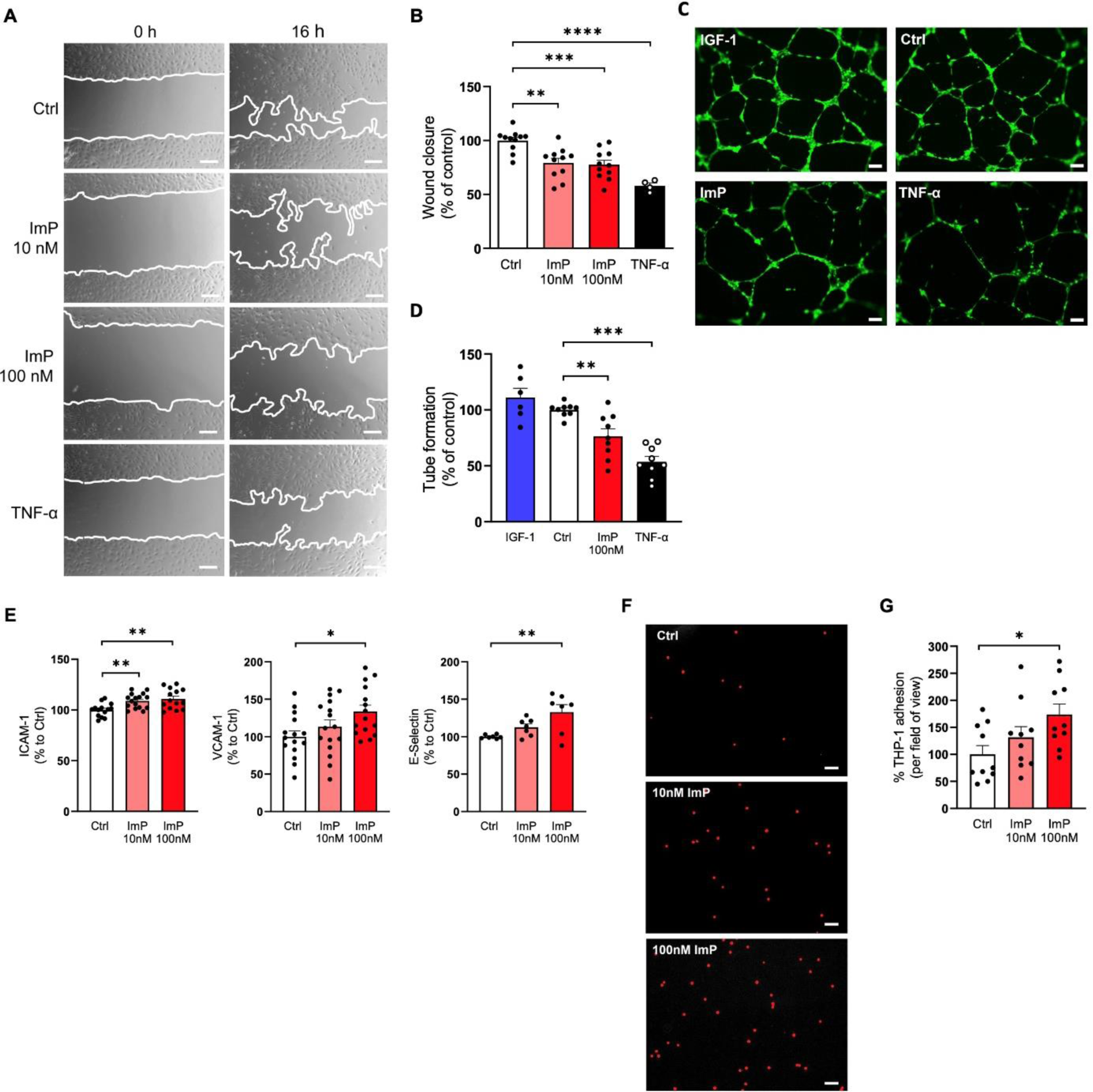
ImP impairs migratory and angiogenic properties, and promotes inflammatory activation of endothelial cells. **A,** Scratch wound healing assay. Representative migration images of HAECs treated with and without ImP or TNF-α for 16 h (10X magnification, scale bars represent 200 µm). **B,** Quantification of migration capacity (*n* = 4 for TNF-α group and *n* = 11 for all other groups). Comparison of wound healing area between control (Ctrl) and treatment with 10 nM ImP, 100 nM ImP or 10 ng/ml TNF-α. **C,** Matrigel tube formation assay. Representative images of endothelial tube formation upon 16 h treatment with IGF-1 (50 ng/ml), control (Ctrl), ImP (100 nM) and TNF-α (10 ng/ml) were taken after Calcein-AM staining (green) (10X magnification, scale bars represent 100 µm). **D,** Endothelial tube formation was quantified by the total number of segments as percentages (*n* = 6 for IGF-1 group and *n* = 9 for all other groups). **E,** ImP increases expression of pro-inflammatory cellular adhesion molecules ICAM-1, VCAM-1 and E-Selectin. Quantitative data are shown as the percentage of gated mean fluorescence intensity (MFI) to control (*n* = 15 for ICAM-1 and VCAM-1 analysis, *n* = 15 for E-Selectin analysis). **F,** Representative images of DiI-labeled THP-1 monocytes (red dots) adhered to endothelial cells upon 24 h stimulation with 10 nM and 100 nM ImP (10X magnification, scale bars represent 100 µm). **G,** Quantification of THP-1 monocyte adhesion to HAECs per field of view in percentage (*n* = 10 per group). Data are shown as mean ± SEM and were calculated by one-way ANOVA followed by the Bonferroni’s post hoc analysis for multiple comparisons (B, D and E)) or by Kruskal-Wallis test followed by the Dunn post hoc analysis for multiple comparisons (G). *\*P* < 0.05, *\*\*P* < 0.01, *\*\*\*P* < 0.001, *\*\*\*\*P* < 0.0001.

### Imidazole propionate promotes endothelial inflammation by inducing the expression of adhesion molecules

Since atherosclerotic lesions in ImP treated *Apoe^-/-^*mice displayed an inflammatory phenotype (containing increased numbers of CD68^+^ macrophages), we next evaluated whether impaired EC functions were accompanied by pro-inflammatory activation. To this end, we performed FACS analyses to determine the expression of endothelial cell adhesion molecules on HAECs as markers of endothelial inflammation and observed a dose-dependent increase in the expression of intercellular adhesion molecule 1 (ICAM-1), E-selectin and vascular cell adhesion molecule-1 (VCAM-1) (Figure 3E). The increased expression of adhesion molecules on endothelial cells following ImP treatment translated to increased monocyte adhesion (Figures 3F and 3G). Since activated endothelium releases soluble VCAM-1 (sVCAM-1) leading to increased circulatory sVCAM-1 levels in blood,^24^ we also evaluated the interaction between circulating sVCAM-1 and ImP levels in a second cohort (the GeneBank cohort) consisting of consecutive patients undergoing elective diagnostic coronary angiography or elective cardiac computed tomographic angiography. Soluble VCAM-1 significantly correlated with circulating ImP levels (Figure S2) further supporting pro-inflammatory actions of ImP leading to increased expression of adhesion molecules that may explain increased monocyte levels in the plaques.

### Imidazole propionate impairs arterial regeneration after carotid injury

In view of the observed detrimental effects of ImP on EC functions *in vitro*, we evaluated the systemic relevance of ImP-mediated effects on vascular regeneration using a mouse carotid injury model (Figure 4A). To this end, mice were subjected to ImP or control vehicle administration for three weeks before undergoing carotid artery injury. Three days after injury, vascular regeneration was evaluated by Evans blue staining, which determines the denuded area (blue-stained). Our results show that treatment with ImP significantly impaired the wound healing process as compared to control animals (Figures 4B and 4C) and, thus, highlight the deleterious effects of ImP on vascular repair mechanisms *in vivo*.

**Figure 4:**
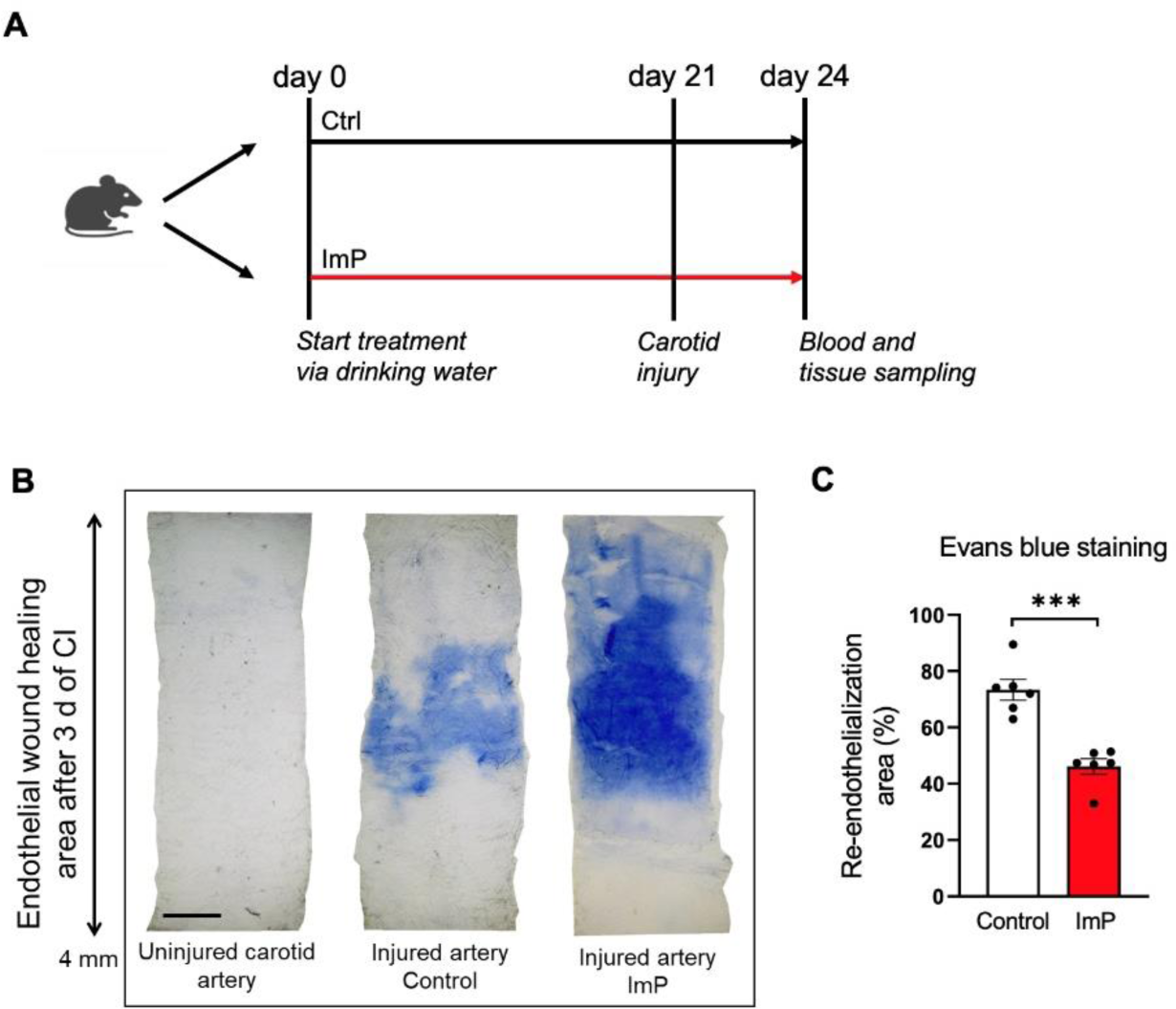
ImP impairs impairs arterial regeneration after carotid injury *in vivo*. **A,** Schematic illustration of the experimental setup for the murine carotid injury model. Control or ImP was administered to adult C57BL/6J mice in drinking water for a total of 24 days. At day 21, carotid injury (CI) was induced to the left carotid artery and vascular re-endothelialization was analyzed after a healing period of three days by Evans blue staining. **B,** Representative *en face* images of Evans blue-stained carotid arteries: uninjured carotid artery, injured carotid artery of control and ImP group. The blue-stained area corresponds to the denuded area of injured carotid arteries (5X magnification, scale bar represents 500 µm). **C,** Quantification of re-endothelialization as the ratio of blue-stained area to total injured area in percentage. Data are shown as mean ± SEM and were calculated by unpaired two-tailed Student’s *t*-test between control and ImP group (*n* = 6 per group). *\*\*\*P* < 0.001.

### Imidazole propionate suppresses PI3K/AKT signaling

To elucidate the underlying molecular mechanism, RNA sequencing of ImP-treated and untreated HAECs was performed. A total number of 52 genes were differentially expressed between the ImP and control conditions, of which 18 genes were upregulated and 34 genes were downregulated in ImP-treated ECs compared to control (Figures 5A and 5B). Notably, several genes involved in angiogenesis were differentially expressed after ImP treatment, including *ADGRL2*, *CALCRL*, *FLT1*, *TM4SF18*, *DCHS1*, *FLRT2*, *ANGPTL4*, *ARGHGAP22* and *IL32* (Figure S3). For example, Latrophilin 2 (*LPHN2*) receptor, also known as *ADGRL2,* was downregulated in response to ImP stimulation. This gene is linked to angiogenic and neurotrophic effects in ECs and mouse tissue explants through de-glycosylated leucine-rich-2-glycoprotein 1 (LRG1)/LPHN2 complex.^25^ Other pro-angiogenic genes downregulated by ImP were Calcitonin receptor-like (*CALCRL*), a G protein-coupled receptor,^26^ vascular endothelial growth factor receptor 1 (*FLT1*),^26,27^ and transmembrane 4 L six family 18 (*TM4SF18*)^27^ known to promote endothelial migration and proliferation *in vitro*. The expression of two genes involved in heart development and morphogenesis regulating endothelial cell migration was also decreased by ImP: *DCHS1*, a member of the cadherin superfamily^28^, and *FLRT2*, a member of the fibronectin leucine-rich transmembrane family. In contrast, the expression of angiogenesis-inhibitory genes, including angiopoietin-like 4 (*ANGPTL4*),^29^ a Rac-specific GTPase isoform of *ARHGAP22* (p68RacGAP)^30^ and interleukin 32 (*IL-32*),^31^ were upregulated by ImP. Thus, the gene expression pattern may shed light on the consistent reduction in endothelial wound healing capacity following ImP treatment.

**Figure 5:**
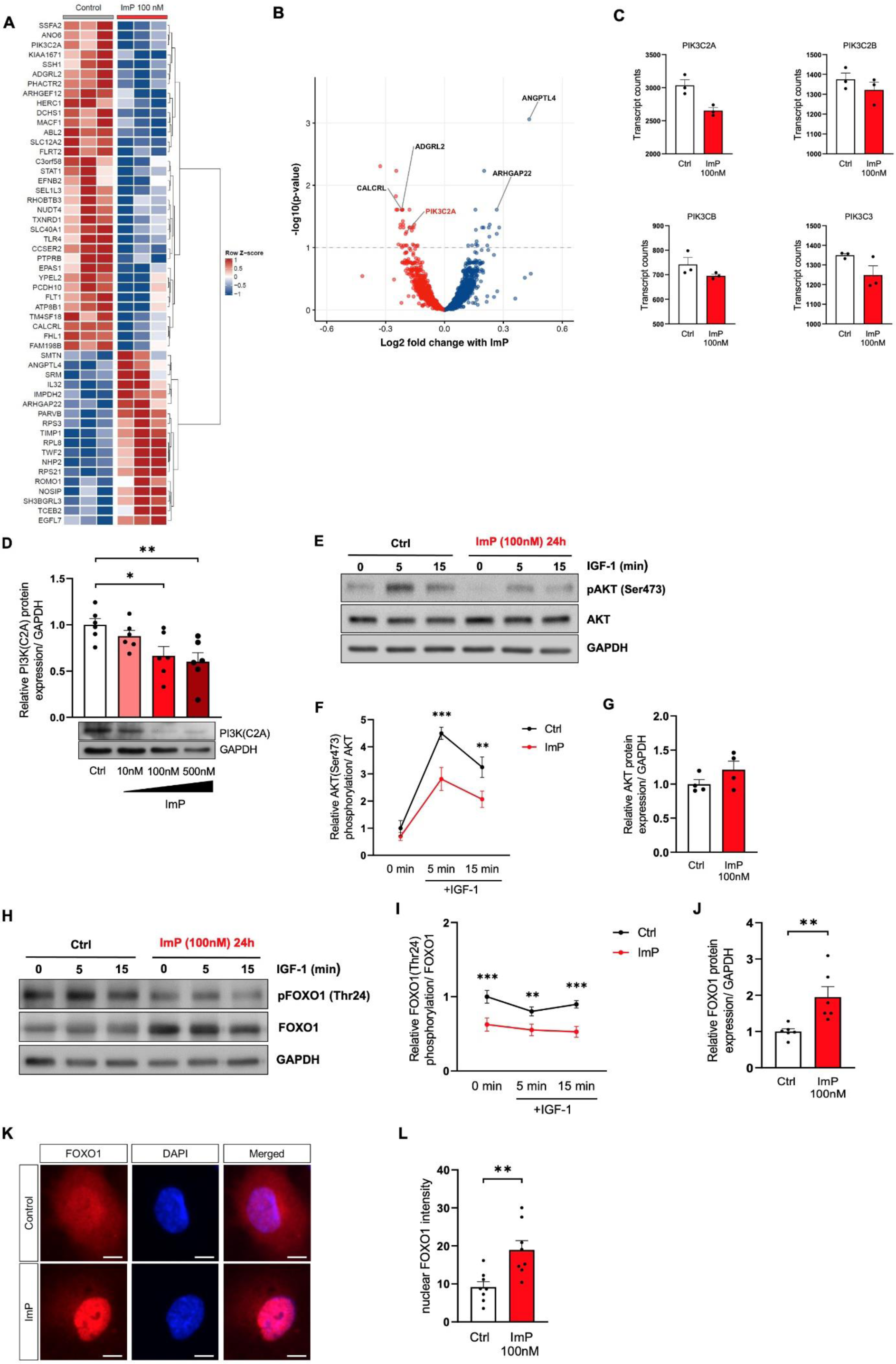
Imidazole propionate suppresses PI3K/AKT signaling and promotes nuclear translocation of FOXO1 in endothelial cells. RNA sequencing analysis of HAECs and differentially expressed genes are illustrated in a heatmap and volcano plot: **A,** Heatmap with 52 differentially expressed genes between ImP and control conditions (adjusted *P*-value < 0.1). Gene counts were transformed to Z-scores (i.e. zero mean, unit variance scaling). The dendrogram on the right was drawn using hierarchical clustering (Ward’s method). **B,** Volcano plot. Red dots represent genes with lower expression in ImP-treated HAECs compared to control group, while blue dots represent genes with higher expression. X-axis denotes the log2 fold change values and the Y-axis shows the –log10 adjusted *P*-values. **C,** Transcript counts from genes of phosphoinositide 3-kinase (PI3K) family of different isoforms including *PIK3C2A*, *PIK3C2B, PIK3CB* and *PIK3C3*. Quantitative data are from three independent experiments (*n* = 3). **D,** Dose-dependent quantification of PI3K(C2A) protein levels in endothelial cells stimulated with control medium, 10 nM ImP, 100 nM ImP or 500 nM ImP. Below are representative western blots of PI3K(C2A) expression normalized to GAPDH (*n* = 6 per group). **E,** Endothelial cells were treated with 100 nM ImP for 24 h followed by IGF-1 stimulation for 0, 5 and 15 min. Representative western blots show protein levels of phospho-AKT(Ser473), AKT and GAPDH (*n* = 4 per group). **F,** Quantification of relative AKT(Ser473) phosphorylation normalized to total AKT protein expression. **G,** Quantification of relative total AKT protein levels normalized to GAPDH expression. **H,** Representative western blots illustrate the protein expression of phospho-FOXO1(Thr24), FOXO1 and GAPDH expression (*n* = 6 per group). **I,** Quantification of relative FOXO1(Thr24) phosphorylation normalized to total FOXO1 protein expression. **J,** Quantification of relative FOXO1 protein level normalized to GAPDH expression. **K,** Immunostaining of endothelial cells showing nuclear sublocalization of FOXO1 (red) and DAPI-stained nucleus (blue) (40X magnification, scale bars represent 10 µm). HAECs were treated with and without 100 nM ImP for 24h. (L) Quantification of nuclear FOXO1 intensity between control and ImP (*n* = 8 per group). Data are shown as mean ± SEM were calculated by one-way ANOVA followed by the Bonferroni’s post hoc analysis (D), two-way ANOVA followed by the Bonferroni’s post hoc analysis for multiple comparisons between control and ImP at different time points (F and I), or unpaired two-tailed Student’s *t*-test between two groups (G, J and L). *\*P* < 0.05, *\*\*P* < 0.01, *\*\*\*P* < 0.001.

Moreover, we found reduced expression of *PIK3C2A,* which encodes phosphatidylinositol-4-phosphate 3-kinase catalytic subunit type 2 (Figure 5C), an enzyme that belongs to the phosphoinositide 3-kinase (PI3K) family^32^ known to be activated upon insulin receptor stimulation.^33^ Western blot analysis confirmed reduced expression of PI3 Kinase Class II α (PI3K(C2A)) also on protein level (Figure 5D). To further explore whether ImP modulates signaling pathways linked to PI3K, we pre-treated ECs with ImP for 24 h followed by insulin receptor activation with IGF-1 stimulation. We observed that ImP reduced AKT phosphorylation (at the activation site of Ser473) (Figures 5E to 5G), and increased FOXO1 transcriptionally activity. Since the forkhead box O transcription factors (FOXOs) are established effectors of the PI3K/AKT pathway and essential regulators of EC growth,^34^ we sought to examine whether dysregulation of FOXO family members links ImP-induced suppression of the PI3K/AKT pathway with impaired angiogenic activity in ECs. FOXO1 is the predominant FOXO family member in endothelial cells.^35^ Activation of the PI3K pathway, e.g., upon insulin receptor stimulation, inhibits FOXO1 through AKT-dependent phosphorylation, leading to inhibition of DNA binding, nuclear exclusion, and subsequent sequestration in the cytoplasm.^35^ Nuclear exclusion of FOXO1 leads to activation of a number of angiogenesis-promoting genes in ECs, thereby promoting endothelial angiogenic activity.^34^ In HAECs, ImP decreased phosphorylation of FOXO1 at AKT-dependent site and led to FOXO1 nuclear translocation (Figures 5H to 5L).

Of note, siRNA-mediated FOXO1 knockdown abolished the ImP-induced increase in ICAM-1 and VCAM-1 levels as observed to control HAECs (Figures 6A and 6B). Consistent with these observations, we found improved vascular regeneration after carotid injury in ImP-treated EC-specific *FOXO1* knockout mice (*Cdh5-CreERT2*(+/–)*/FOXO1^fl/fl^*) compared to control littermates without Cre recombinase activity (*Cdh5-CreERT2*(+/+)*/FOXO1^fl/fl^*) (Figures 6C to 6E). Together, these findings support the involvement of a FOXO1-dependent pathway in ImP signaling leading to reduced vascular regeneration and a pro-inflammatory response.

**Figure 6:**
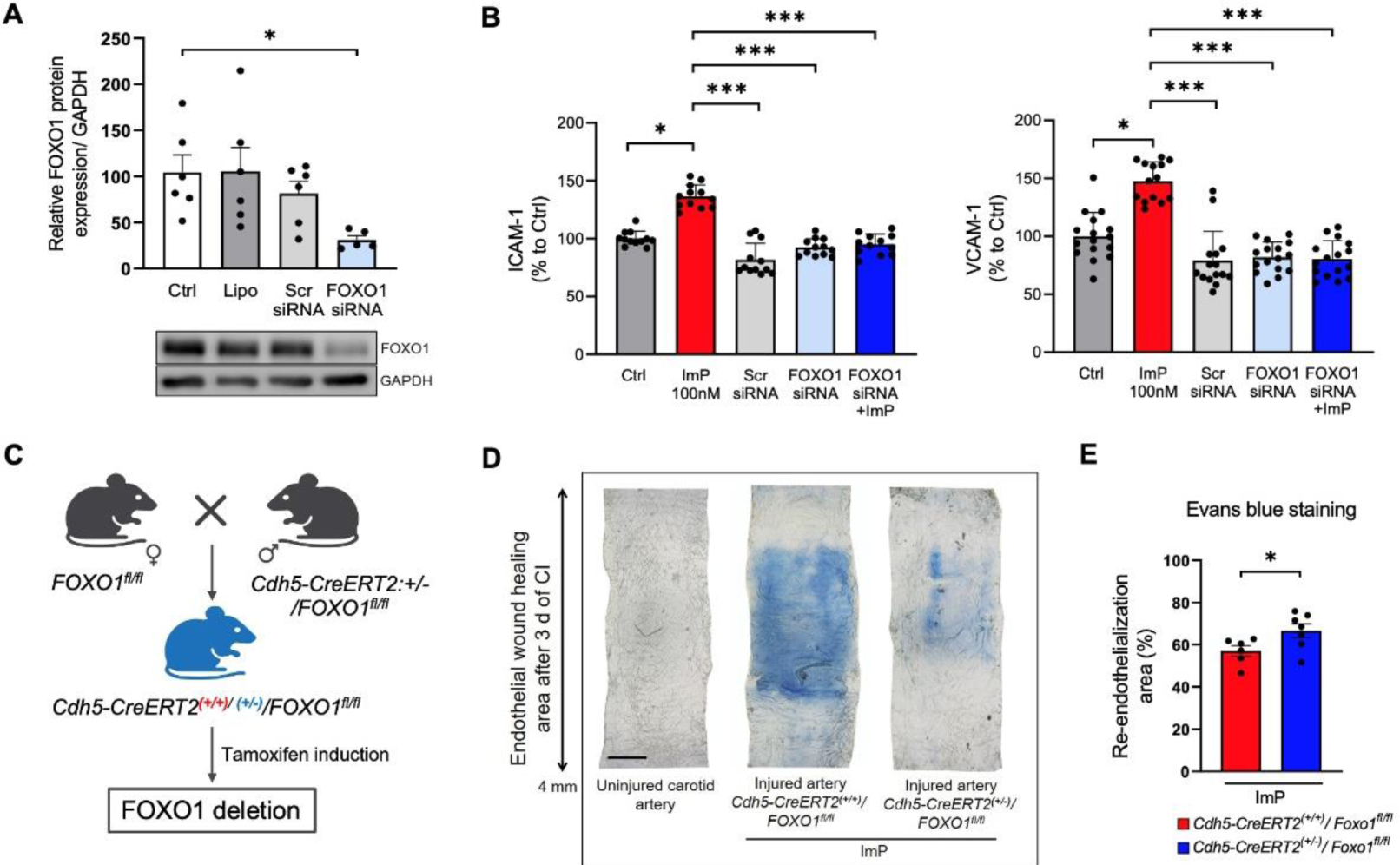
Inhibition of endothelial-specific FOXO1 restores ImP-mediated effects on endothelial inflammation and vascular damage. **A,** Quantification of FOXO1 knockdown in endothelial cells on protein levels. Below are the representative western blots of FOXO1 protein levels after 4h of siRNA transfection. Total FOXO1 protein expression was normalized to GAPDH expression (*n* = 5 for FOXO1 siRNA group and *n* = 6 for all other groups). **B,** Flow cytometric analysis representing data of cellular adhesion surface expression of ICAM-1 (*n* = 12) and VCAM-1 (n = 16) in FOXO1-silenced HAECs followed by ImP treatment for 24 h. Control groups include control medium, scramble (Scr) siRNA and FOXO1 siRNA. **C,** Schematic illustration of endothelial-specific FOXO1 knockout mice using tamoxifen-inducible Cre-loxP system. *LoxP*-flanked FOXO1 mice (*FOXO1^fl/fl^*) were crossbred with transgenic mice expressing the tamoxifen-inducible vascular endothelial-specific cadherin (*Cdh5*) promoter-driven CreERT2 recombinase (*Cdh5-CreERT2:+/-*). Deletion of FOXO1 in endothelial cells was induced in the presence of Cre recombinase through intraperitoneal injections of tamoxifen (100 mg/kg per body weight) once a day for 5 consecutive days. FOXO1 knockout mice are referred to as *Cdh5-CreERT2*(+/–)*/FOXO1^fl/fl^*, while control animals were littermates without Cre recombinase activity and are referred to as *Cdh5-CreERT2*(+/+)*/FOXO1^fl/fl^*. **D,** Representative *en face* images of Evans blue-stained carotid arteries: uninjured carotid artery, injured carotid artery from control littermate (*Cdh5-CreERT2*(+/+)*/FOXO1^fl/fl^*) and from FOXO1 deficiency mice (*Cdh5-CreERT2*(+/–)*/FOXO1^fl/fl^*). The blue-stained area corresponds to the denuded area of injured carotid arteries (5X magnification, scale bar represents 500 µm). **E,** Quantification of re-endothelialization as the ratio of blue-stained area to total injured area in percentage (*n* = 6 per group). Data are shown as mean ± SEM and were calculated by one-way ANOVA followed by the Bonferroni’s post hoc analysis (A), Kruskal-Wallis test followed by the Dunn post hoc analysis for multiple (B) or unpaired two-tailed Student’s *t*-test between control littermates without Cre recombinase activity and FOXO1 deficiency animals upon ImP treatment (E). *\*P* < 0.05, *\*\*\*P* < 0.001.

## DISCUSSION

Our study demonstrates that plasma levels of the gut microbial metabolite ImP is associated with increased risk for prevalent CAD in humans even after adjusting for traditional cardiovascular risk factors. Furthermore, we provide evidence that ImP causally contributes to the development of atherosclerosis by impairing EC functions in both *in vitro* and *in vivo* models. We identified a potential mechanism, whereby ImP reduces endothelial PI3K/AKT signalling, thereby limiting their proliferative and migratory activities. Finally, the present results show that ImP impairs endothelial regeneration after arterial injury and accelerates the progression of atherosclerosis (Figure 7).

**Figure 7:**
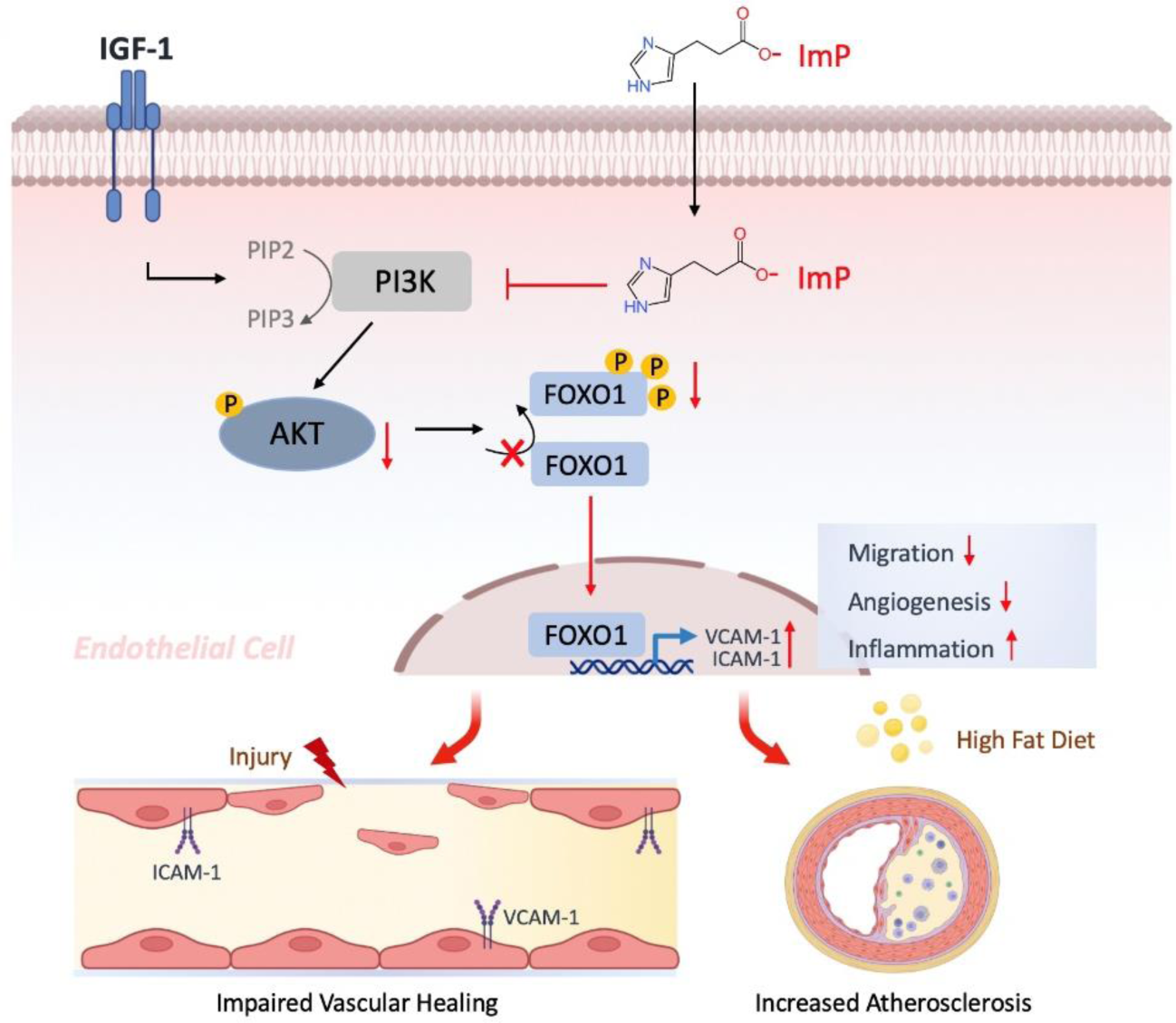
Novel cellular mechanism of Imidazole propionate (ImP) involved in endothelial dysregulation. The PI3K/AKT signaling pathway regulates the transcriptional activity of FOXO1 in endothelial cells. Upon IGF-1 stimulation, PI3K is activated through the generation of phosphatidylinositol 4,5-bisphosphate (PIP2) to phosphatidylinositol (3,4,5)-trisphosphate (PIP3), thus, resulting in subsequently AKT activation that directly phosphorylates and thereby inactivates FOXO1. AKT-mediated phosphorylation of FOXO1 induces FOXO1 nuclear exclusion and inhibits FOXO1 transcriptional activity. Upon treatment with Imidazole propionate (ImP), PI3K/AKT signaling is significantly downregulated leading to decreased phosphorylation and increased nuclear accumulation of FOXO1, thus, leading to the expression of FOXO1 downstream targets ICAM-1 and VCAM-1 in endothelial cells. Combined, these actions promote endothelial cell inflammation, impaired vascular healing process and accelerates the progression of atherosclerosis development (created with BioRender.com).

Accelerated vascular disease is a major determinant of increased morbidity and mortality risk in patients with diabetes mellitus^36^ and injury to the arterial wall is considered as a crucial step in initiation and progression of atherosclerotic vascular disease.^1^ Endothelial denudation upon arterial wall injury induces a cascade of cellular and molecular actions, ultimately leading to vascular wall remodeling, which initiates or promotes the development of atherosclerotic vascular disease.^37^ This process includes leukocyte chemotaxis with secretion of an array of cytokines and growth factors that govern the migration and proliferation of smooth muscle cells and interstitial collagen gene expression leading to extracellular matrix deposition following injury.^38^ While proper endothelial cell repair may mitigate pathologic alterations of the vascular wall, impaired regenerative capacity of the endothelium facilitates adverse vascular wall remodeling and thus, promotes atherosclerosis.^39,40^ A dysfunctional endothelium is characterized by an impairment of fundamental cellular properties such as migration, proliferation and sprout formation.^41^ Moreover, it is associated with inflammatory activation.^42^

Despite enormous efforts to elucidate mechanisms of EC dysfunction over the past decades, contributing pathways at the systems level still remain incompletely understood. One hypothesis is that defective insulin receptor signaling, e.g., due to impaired glucose tolerance or diabetes,^3^ is a major cause of EC dysfunction and regenerative potential and is associated with substantially increased risk of ACVD.^43^

In recent years, increasing evidence demonstrated that alterations of the gut microbiota, its metabolic capacities, intestinal gene expression, and intestinal immune system are important contributing factors to cardiometabolic diseases,^11–13,44^ which may contribute to endothelial dysfunction and ACVD.^45^ The altered microbiota associated with cardiometabolic diseases can produce diseases causing and modifying metabolites.^46^ ImP is increased in the portal and peripheral blood of obese patients with versus without type 2 diabetes.^15^ These findings have been confirmed in patients of different geographic location and ImP was found to be associated with low bacterial gene richness.^16^ Recently, ImP was found to be increased in patients with heart failure.^18^ However, it is unknown how ImP affects endothelial cell function and ACVD. Our findings demonstrate a link between ImP levels and increased risk for prevalent CAD in humans and we demonstrated causality by chronic administration of ImP to atherosclerosis prone mice in a lipid independent mechanism. Rather, our *in vitro* studies suggest deleterious effects of ImP on endothelial cells reducing the regenerative potential of the endothelium after vascular injury. In particular, we demonstrate that ImP reduces the ability of endothelial cells to activate insulin receptor-induced PI3K/AKT signaling by downregulating the PI3K(C2A). The impairment of PI3K/AKT signaling is accompanied by decreased phosphorylation and increased nuclear accumulation of FOXO1, thereby attenuating the repair capacity of the endothelium. These findings are consistent with previous reports showing that FOXO1 decelerates metabolic activity by reducing glycolysis and mitochondrial respiration.^34^ However, endothelial FOXO1 deficiency contributed to the reversal of the deleterious effects induced by ImP on endothelial cell physiology, potentially influencing inflammatory response in ECs and vascular repair mechanisms positively. Taken together, our findings demonstrate a link between ImP and endothelial cell dysfunction, which involves a deregulated PI3K/AKT/FOXO1 signaling axis. Moreover, they demonstrate a causative role of ImP in the development of ACVD, and thus suggest therapeutic effects of targeting ImP-producing gut microbial pathways in cardiovascular disease prevention.

## Clinical perspective: What is new?

- Circulatory imidazole propionate (ImP) levels are higher in patients with atherosclerotic coronary artery disease (CAD) than in patients without CAD.
- ImP promotes development of atherosclerosis in *Apoe^-/-^* mice.
- ImP attenuates vascular repair capacity after injury in mice.
- ImP impairs insulin receptor signaling in endothelial cells by suppressing PI3K/AKT pathway and subsequent activation of FOXO1 transcription factor.

## What are the clinical implications?

- ImP is causally linked to atherosclerotic cardiovascular disease.
- Targeting ImP-producing gut microbial pathways may provide a novel concept in cardiovascular disease prevention.

## Nonstandard abbreviations and acronyms

ACS: acute coronary syndromes
ACVD: atherosclerotic cardiovascular disease
Apoe^-/-^: apolipoprotein E knockout
CAD: coronary artery disease
Cdh5: vascular endothelial-specific cadherin
CI: carotid injury
EC: endothelial cell
FOXO1: forkhead box O transcription factor 1
HAECs: human aortic endothelial cells
ImP: Imidazole propionate
LCA: left carotid artery
LDL: low-density lipoprotein
PI3K: phosphoinositide 3-kinase
PIK3C2A: phosphatidylinositol-4-phosphate 3-kinase catalytic subunit type 2
sVCAM-1: soluble VCAM-1
TC: total cholesterol
UHPLC: ultra high-performance liquid chromatography
VLDL: very low-density lipoprotein

## Acknowledgements

The authors would like to thank M. Moobed and N. Rösener for excellent assistance on cell culture experiments, and Y. Jansen and S. Bayasgalan on performing lipid profiling *in vivo*.

## Author contributions

Authors contributed equally as senior authors: Ulf Landmesser, Fredrik Bäckhed and Arash Haghikia.

V.N, designed and performed the *in vitro* and *in vivo* experiments, and contributed to the bioinformatic data analysis. A.C., K.R.B, M.S. and F.B. designed the ImP drinking protocol and performed mass spectrometry analysis in human and mouse samples. M.K., I.D., D.M.L., U.L. and A.H. coordinated the LipidCardio study and contributed clinical study samples. J.S. contributed to the bioinformatic data analysis of the human study. L.R., P.R.R, E.T.S. and N.K. provided general experimental support. B.V. performed RNA-seq bioinformatics analysis. Y.D. and C.W. performed the lipid analysis. J.L. and M.P. provided endothelial-specific FOXO1 knockout animals and supported experimental work. M.N., P.K. and E.S.T. supported research and provided valuable input. M.F. and S.L.H. performed the mass spectrometry analysis in human samples and ELISA of soluble VCAM-1. A.H., F.B. and U.L. supervised the project and contributed equally. V.N. and A.H. wrote the paper. All authors reviewed and approved the manuscript.

## Supplementary Data

**Figure S1:**
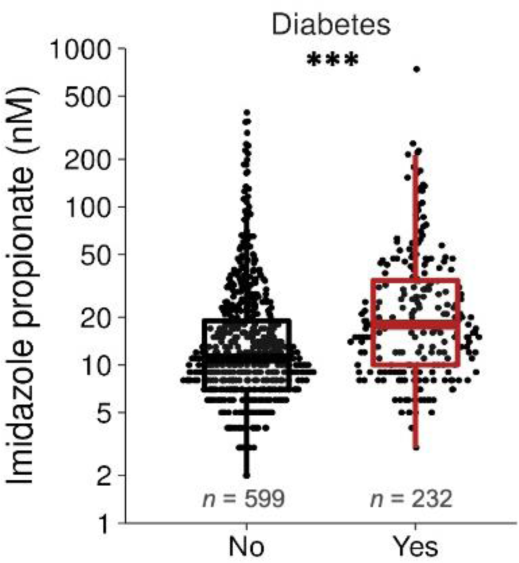
Imidazole Propionate is increased in patients with type 2 diabetes. Graph shows the comparison of ImP levels between patients without and with type 2 diabetes (*n* = 831, *\*\*\*P* < 0.001). Data is shown as boxplots demonstrating the middle line as the median, the lower and upper hinges as the first and third quartiles, and the whiskers represent 10th and 90th percentiles; *P*-values were analyzed using Wilcoxon rank sum test.

**Figure S2:**
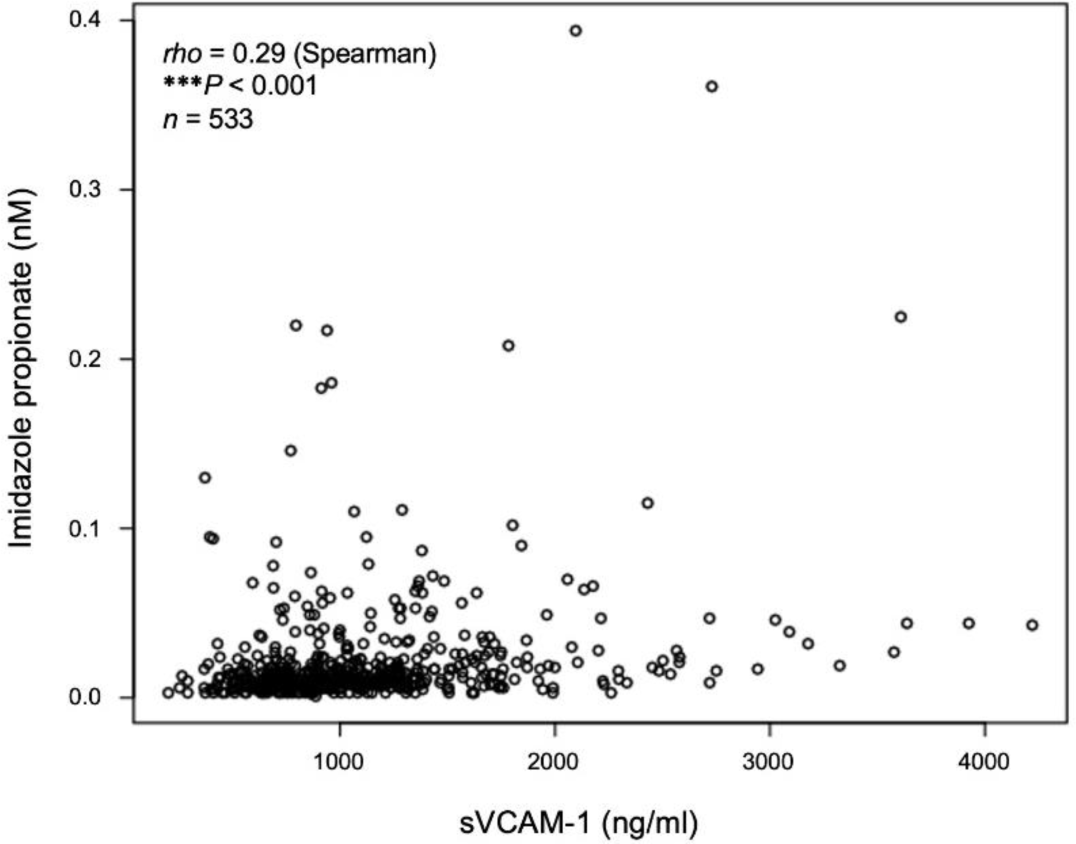
Imidazole propionate significantly correlates with increased circulatory soluble VCAM-1 levels in blood of patients. Graph demonstrates the Spearman correlation between plasma levels of ImP (nM) and soluble VCAM-1 (ng/ml) from the GeneBank study supporting a putative link between ImP and VCAM-1 levels (*n* = 533, *\*\*\*P* < 0.001, Spearmańs *rho* = 0.29).

**Figure S3:**
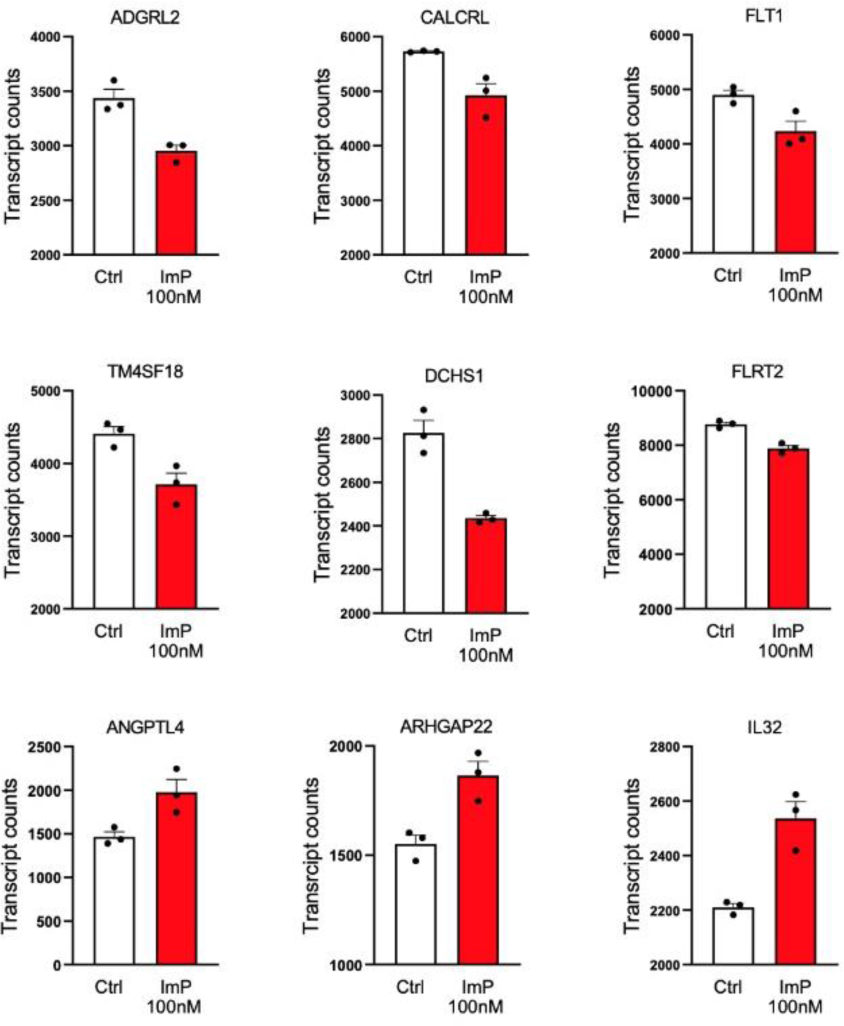
Imidazole Propionate downregulates a number of angiogenic genes in endothelial cells. Box plot charts illustrating key down- or upregulated genes from the RNA sequencing analysis involved in angiogenesis pathway including *ADGRL2, CALCRL, FLT1, TM4SF18, DCHS1, FLRT2, ANGPTL4, ARGHGAP22,* and *IL32* between the control and ImP group. Represented data are transcript counts differentially expressed in endothelial cells after treatment with and without ImP (100 nM) and from three independent experiments (*n* = 3).

**Table S1:** Experimental models.

| Cell lines and animals/strains. | Source | Identifier |
| --- | --- | --- |
| Human aortic endothelial cells (HAECs) | Cell Applications | 304-05a |
| THP-1 monocytes | N/A | N/A |
| C57BL/6J mice (male) | Charles River Laboratories | N/A |
| *Apoe*<sup>-/-</sup> mice (male) (B6.129P2) | Charles River Laboratories | N/A |
| *Cdh5-CreERT2*<sup>(+/-)</sup>/*FOXO1*<sup>fl/fl</sup> mice<br>*Cdh5-CreERT2*<sup>(+/+)</sup>/*FOXO1*<sup>fl/fl</sup> mice (male and female) | Max Delbrück Center (MDC; Berlin, Germany) | N/A |

**Table S2:**
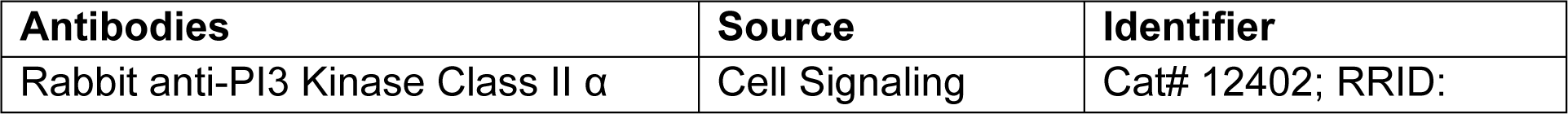
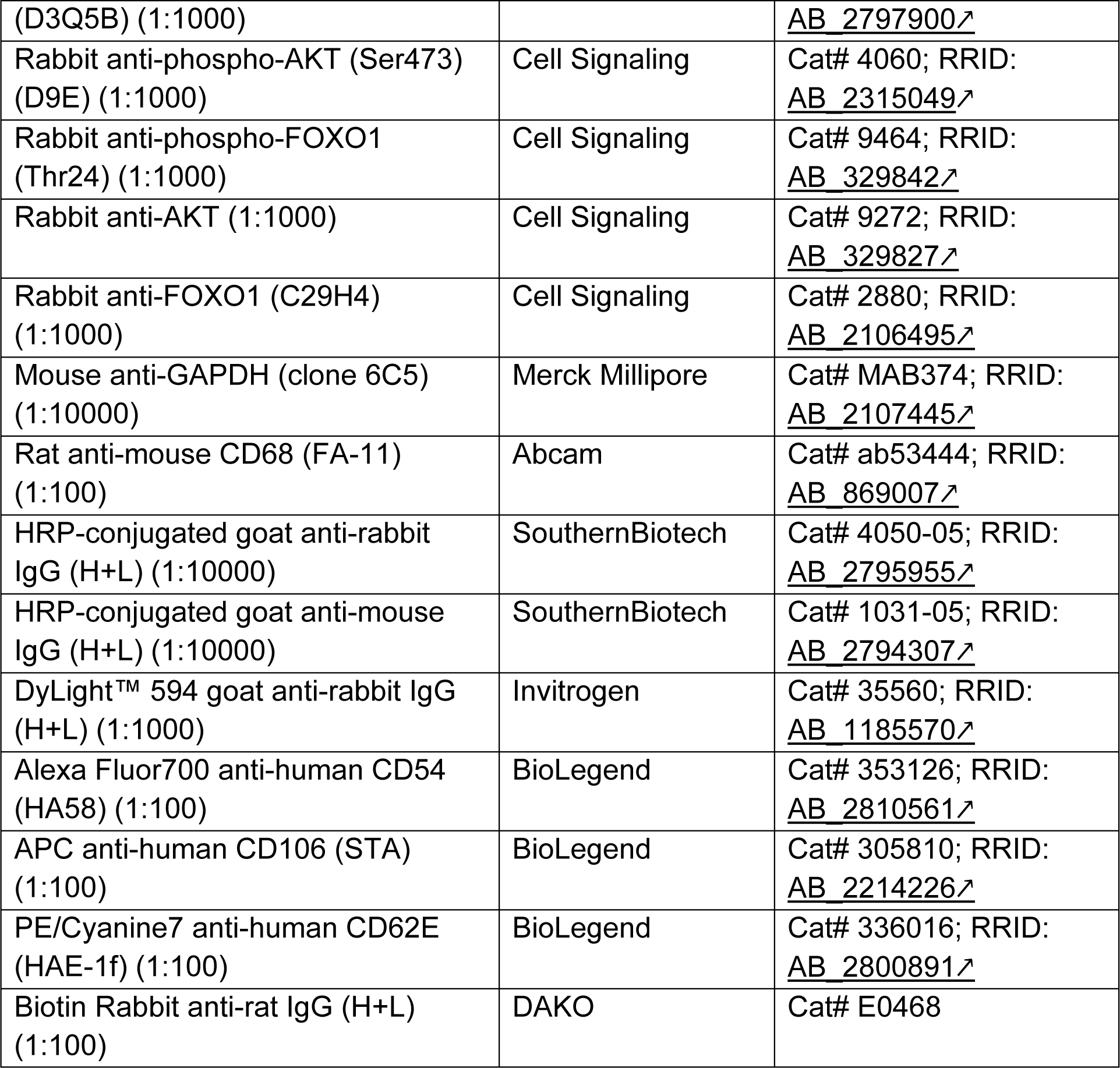
Antibody reagents and resource.

**Table S3:** Oligonucleotides.

| Oligonucleotides | Source | Identifier |
| --- | --- | --- |
| Control siRNA (scrambled) | Santa Cruz | sc-37007 |
| FKHR/FOXO1 siRNA | Santa Cruz | sc-35382 |

**Table S4:**
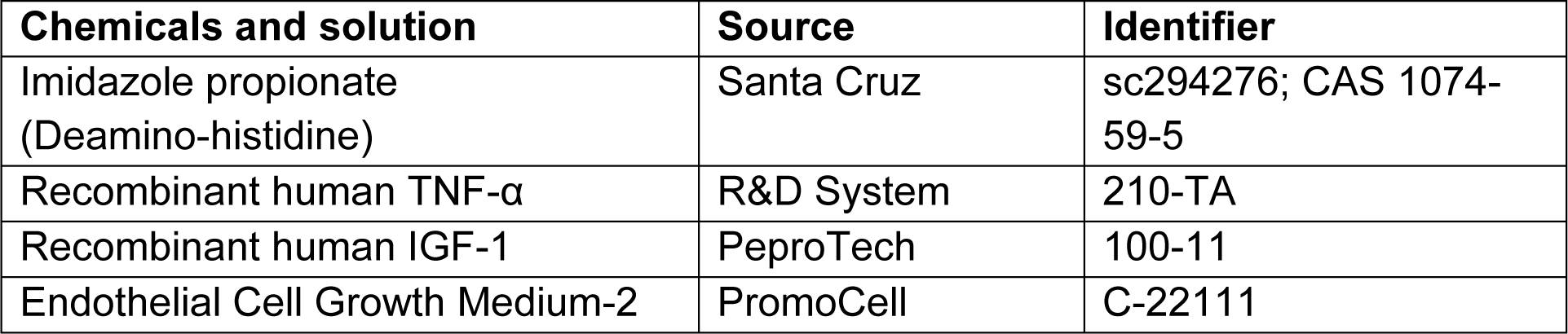
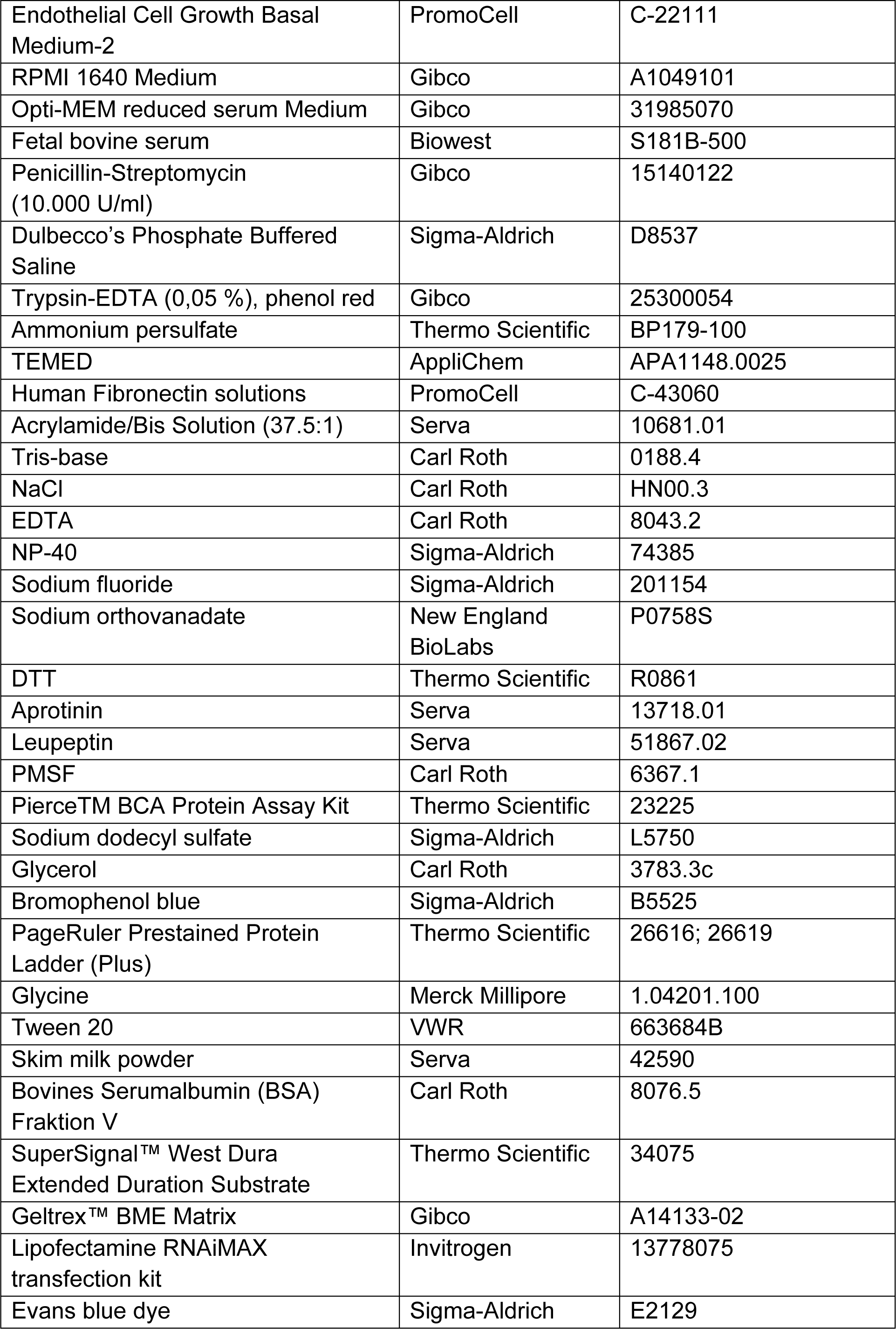
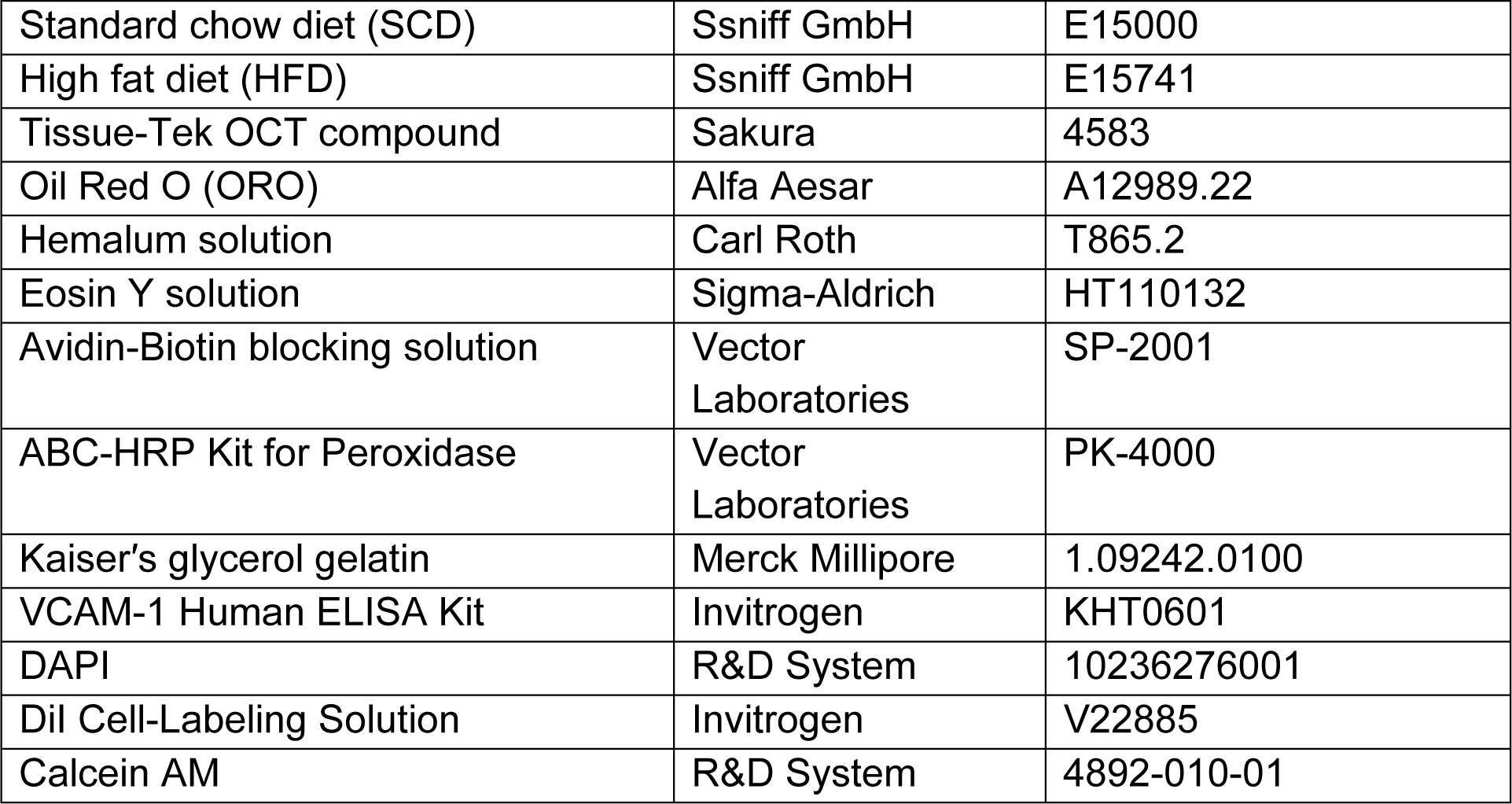
Experimental materials.

